# Nrf2 orchestrates epigenetic regulations and serves as the master regulator of KLF4 expression and activity during arsenic-induced transformation

**DOI:** 10.1101/2024.10.15.617132

**Authors:** Ziwei Wang, Zhuoyue Bi, Jessica Bamrah, Yiran Qiu, Wenxuan Zhang, Haoyan Ji, John D. Haley, Chitra Thakur, Fei Chen

## Abstract

Emerging evidence suggests that Nrf2 plays a pro-carcinogenic role in cancer. Our previous study showed that arsenic-induced Nrf2 activation triggers metabolic reprogramming, leading to the formation of cancer stem-like cells. Here, we further demonstrated that KLF4, a key pluripotency factor, is a direct transcriptional target of Nrf2 in arsenic-exposed human bronchial epithelial cells. ChIP-seq analysis identified multiple Nrf2 binding peaks at the Klf4 gene locus, which overlap with the enhancer markers H3K4me1 and H3K27Ac. Nrf2 knockout reduced both KLF4 expression and enhancer marker enrichment, accompanied by a global decrease in KLF4 binding across the genome. In wild-type (WT) cells, arsenic treatment increased KLF4 binding on genes involved in oncogenic pathways such as STAT3, SOX2, Nrf2, cell growth, Hedgehog, and EMT. We also found that KLF4 engages in a self-feedback loop in response to Nrf2 signaling. Lastly, our data showed that the co-occupancy of Nrf2 and KLF4 is crucial for establishing active enhancer hubs in the genome. These findings suggest that Nrf2’s oncogenic effects are, in part, mediated by Nrf2 dependent self-amplification of KLF4 expression and function. Thus, targeting both Nrf2 and KLF4 could be a promising therapeutic strategy for eliminating cancer stem-like cells.

---

It remains unclear how environmental and other risk factors contribute to the development of human cancers. Aberrant activation of Nrf2 or somatic mutations in Nfe2l2, the gene encoding Nrf2, a cap’n’collar leucine zipper transcription factor, have frequently been detected in several human cancers [1]. Approximately 25 to 30% of human non-small cell lung cancer (NSCLC) are characterized by hyperactive Nrf2, resulting from genomic mutation in Nfe2l2 or its negative regulator, Keap1 [2, 3]. While Nrf2 has primarily been viewed as an oncogenic protein due to its role in promoting the expression of cytoprotective genes in response to oxidative and electrophilic stresses induced by environmental hazards, a few earlier studies have suggested its tumor suppressive activity. In a mouse model of NSCLC, constitutive activation of Nrf2 initiated lung tumor development and promoted early progression from hyperplasia to low-grade tumor, but did not lead to the formation of advanced-grade tumors [4]. In human lung cancer, however, emerging evidence indicates that sustained Nrf2 activation supports metastasis, therapeutical resistance, tumor recurrence, and poor prognosis of cancer patients [5, 6]. It is believed that Nrf2-mediated heme catabolism can cause the accumulation of Bach1, a pro-metastatic protein [7]. Additionally, persistently activated Nrf2 in NSCLCs has been shown to cooperate with CEBPB to remodel enhancers at oncogenic gene loci that are not typically regulated by transiently activated Nrf2 [8]. In pancreatic cancer, Nrf2 is crucial for maintaining cancer growth by upregulating mRNA translation through EGFR and AKT kinase signaling [9].

We previously demonstrated the role of Nrf2 as a master regulator in metabolic reprogramming and the generation of cancer stem-like cells (CSCs) in the human bronchial epithelial cell line BEAS-2B treated with environmentally relevant concentrations of arsenic (As^3+^) [10, 11]. Beyond its known target genes in the antioxidant responses of cells, global ChIP-seq analysis identified a number of genes crucial for the metabolic shift from the mitochondrial citric acid (TCA) cycle to glycolysis, as well as for the shunting pathways of glycolysis, as being regulated by Nrf2. This notion is supported by recent evidence suggesting the regulatory role of Nrf2 in glycolytic metabolism in various cell types, including prostate stem progenitor cells [12], human hepatocarcinoma cells [13], breast cancer cells [14], oral squamous cell carcinoma [15], T cells [16, 17], cardiomyocytes [18], endothelial cells [19], and other cells and tissues [20–22]. A unique feature of Nrf2-mediated glycolytic metabolism in As^3+^-induced transformed or CSCs is the active shunting of glycolytic intermediates into the hexosamine biosynthetic and the serine-glycine pathways [10]. While Nrf2 drives glycolytic metabolism, recent studies also suggest that glycolytic intermediates, such as glyceraldehyde 3-phosphate, can activate Nrf2 by inactivating KEAP1 through S-lactoyl modification [23]. Since KEAP1 is an endogenous inhibitor of Nrf2 that targets it for ubiquitination and proteasomal degradation, this S-lactoyalation of KEAP1 indicates a mutual amplification of Nrf2 activity and glycolytic metabolism.

It is well-established that glycolysis is closely associated with the stemness of the stem cells, progenitor cells, cancer cells, and CSCs [24]. Certain glycolytic metabolites, such as uridine diphosphate N-acetylglucosamine (UDP-GlcNAc) and NADPH, are essential for the activation or function of pluripotency transcription factors. In addition to regulating genes involved in glycolytic metabolism, Nrf2 may directly induce the expression of several genes associated with the stemness of cancer cells. A transient treatment of cells with As^3+^ resulted in a significant enrichment of Nrf2 on pluripotent and/or stem cell marker genes, including Myc, Sox2, Klf4, Cd44, Egfr, Bach1, Fgf1, Ngf, Pdgfb, among others, positioning Nrf2 as a critical promoting factor of CSC formation and functional specialization in carcinogenesis [10, 25]. In the current report, we provide new evidence underscoring the importance of Nrf2 in orchestrating epigenetic regulations on the expression and function of KLF4. Our ChIP-seq analysis revealed multiple Nrf2 binding peaks that overlap with major enhancer markers, H3K4me1 and H3K27Ac, within the KLF4 gene loci. Knocking out Nrf2 reduces the levels of H3K4me1 and H3K27Ac at the KLF4 gene, along with decreased expression and DNA binding of KLF4 in As^3+^-treated cells. These findings, thus, suggest that targeting Nrf2 or its downstream gene, Klf4, could be a novel strategy for cancer therapy.

## Results

### Nrf2 dependency in As^3+^-induced malignant transformation

We previously demonstrated that knocking out (KO) Nrf2 increases the sensitivity of BEAS-2B cells to cytotoxicity when treated with 0.25 μM As^3+^ for 20 days. This led to hypothesis that Nrf2 is essential for As^3+^-induced transformation and the generation of CSCs [25]. To explore this further, we extended As^3+^-treatment to two months. During this prolonged exposure, Nrf2 KO cells, after an initial phase of growth restriction, began proliferating at a rate comparable to wild type (WT) cells in the presence of As^3+^. Both WT and Nrf2 KO cells displayed characteristics of malignant transformation, as indicated by anchorage-independent growth in soft agar. However, while transformed WT cells formed numerous large colonies, transformed Nrf2 KO cells primarily produced small colonies and individual resting cells (Fig. 1A), highlighting the critical role of Nrf2 in cell transformation. Gene expression profiles are typically distinct in transformed or malignant cells. To identify Nrf2-regulated genes during transformation, we performed transcriptomic analysis using RNA-seq on both WT and Nrf2 KO cells, untreated or treated with 2 μM As^3+^ for 6h. We found 3343 genes downregulated in Nrf2 KO cells with a shrunken Log2 FC ≤ 0.25 under control conditions, indicating Nrf2 dependency. An Integrated Cancer Hallmark Gene Set analysis revealed that genes involved in metastasis, evasion of growth suppressors, sustained proliferation, and resistance to cell death were significantly enriched among these Nrf2 dependent genes (Fig. 1B). As^3+^ treatment induced 264 genes in WT cells, particularly in pathways associated with reprogramming energy metabolism and resisting cell death. These changes were not observed in Nrf2 KO cells (Fig. 1B), with key genes such as PLK3, G6PD, FOXO3, ATF4, HK2, PKM, CD44, NGF, LIF, EPAS1, IKBKG, BMP4, GADD45A, among others. These findings are clinically relevant, as the fact that Nrf2 is positively correlated with 604 genes (Rho ≥ 0.5) in a cohort of 1,865 human lung tumors, many of which are Myc target genes or genes are involved in mTORC1, androgen, and NF-κB signaling pathways. These include stemness genes like KLF4, CD44, ITGA6, CD9, and KLF5 (Fig. 1C). Notably, Nrf2 ranked as the top gene positively correlated with KLF4, with a Spearman’s Rho value ≥ 0.5 in human lung cancer (Fig. 1D). Higher expression of Klf4 is linked to poorer overall survival in lung cancer patients with tobacco smoke (Fig. 1E), emphasizing its prognostic relevance.

**Fig.1.**
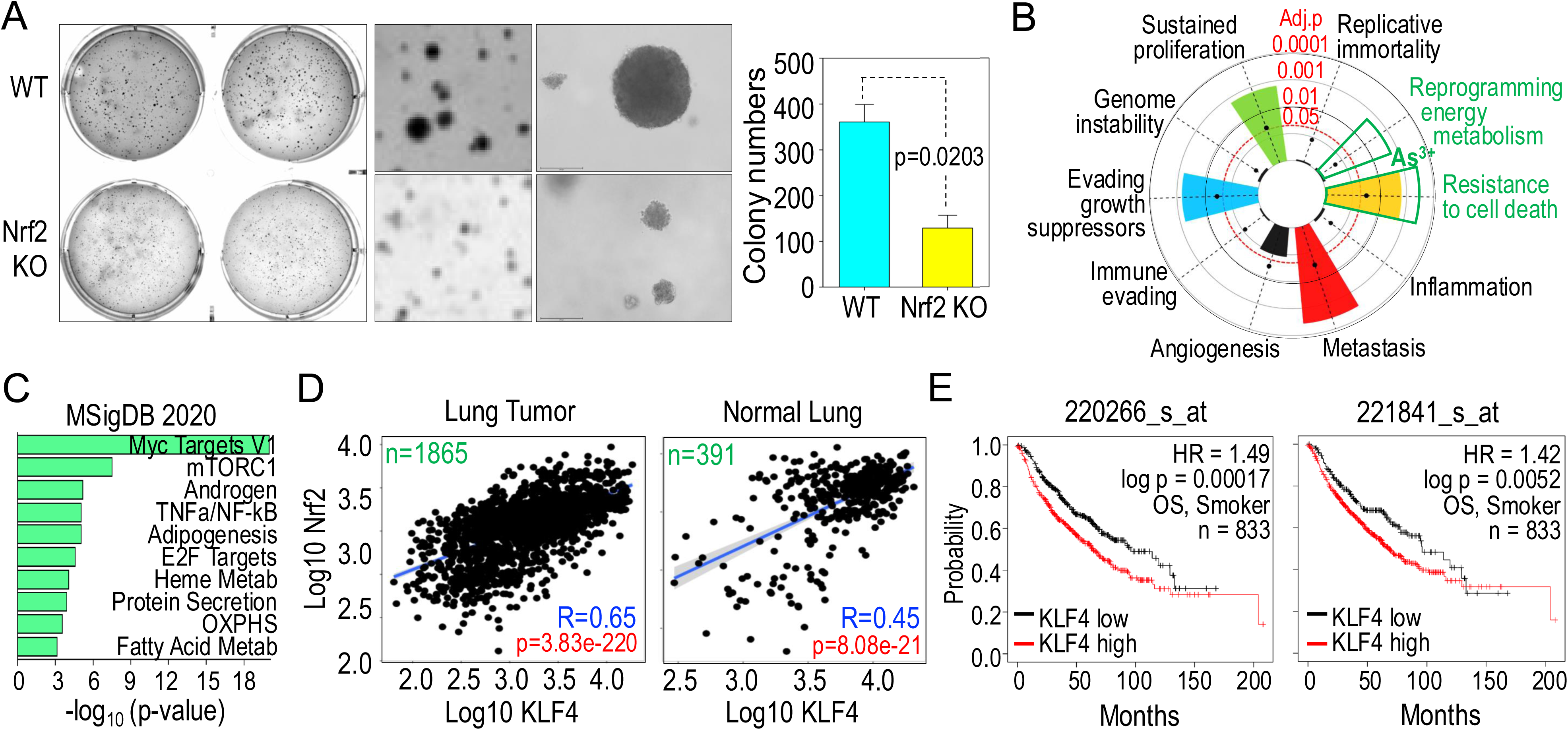
Nrf2 is essential for As^3+^-induced transformation and correlated with KLF4 in lung cancer. **A.** Anchorage-independent growth of the WT and Nrf2 KO cells treated with 0.25 μM As^3+^ for two months in soft agar. Right panel shows quantification of cell colonies of the WT and Nrf2 KO cells with a diameter large than 100 μm in soft agar. **B.** Integrated Cancer Hallmark Gene Set analysis of 3343 genes that showed decreased expression in Nrf2 KO cells as determined by RNA-seq and 264 genes that were induced by 2 μM As^3+^ for 12h in WT cells. **C.** Molecular Signature of Cancer Hallmark pathway analysis of 604 genes that are positively correlated with Nrf2 with a Spearman’s Rho ≥ 0.5 in 1865 human lung tumors. **D.** Nrf2 (Nfe2l2) is the top-one KLF4-correlated genes in human lung cancer and one of the most KLF4-correlated genes in normal human Lung. **E.** High levels of KLF4 expression as determined by two different probes predict poorer over survival of the lung cancer patients associated with tobacco smoke.

### Klf4 is a transcriptional target of Nrf2 that cooperates with enhancers

The strong correlation between Nrf2 and KLF4 in human lung tumors suggests that Nrf2 directly regulates the expression of KLF4. In WT cells, transient treatment with As^3+^ activates Nrf2 and dose-dependently increases the protein levels of KLF4 and MYC, two oncogenic transcription factors associated with stem cells and CSCs. However, this induction of KLF4 is significantly reduced in Nrf2 KO cells (Fig. 2A). Nrf2 ChIP-seq analysis revealed a strong increase in Nrf2 binding induced by As^3+^ within the gene body, as well as upstream and downstream regions of the Klf4 gene (Fig. 2B). These Nrf2 binding clusters, which contain one or two consecutive Nrf2 binding elements known as antioxidant responsive elements (ARE), had not been previously reported. Another notable characteristic of these Nrf2 binding clusters is their colocalization with enhancer markers H3K4me1 and H3K27Ac, designated as e1 to e7 at the Klf4 gene locus (Fig. 2B). The enrichment levels of these enhancer markers are significantly reduced in Nrf2 KO cells. On a genomic scale, our ChIP-seq data indicate a tendency for Nrf2 to colocalize with enhancer markers, suggesting that Nrf2 may coordinate with other epigenic regulators, such as enhancers, to create a permissive chromatin environment for the expression of Klf4 and many other genes.

**Fig. 2.**
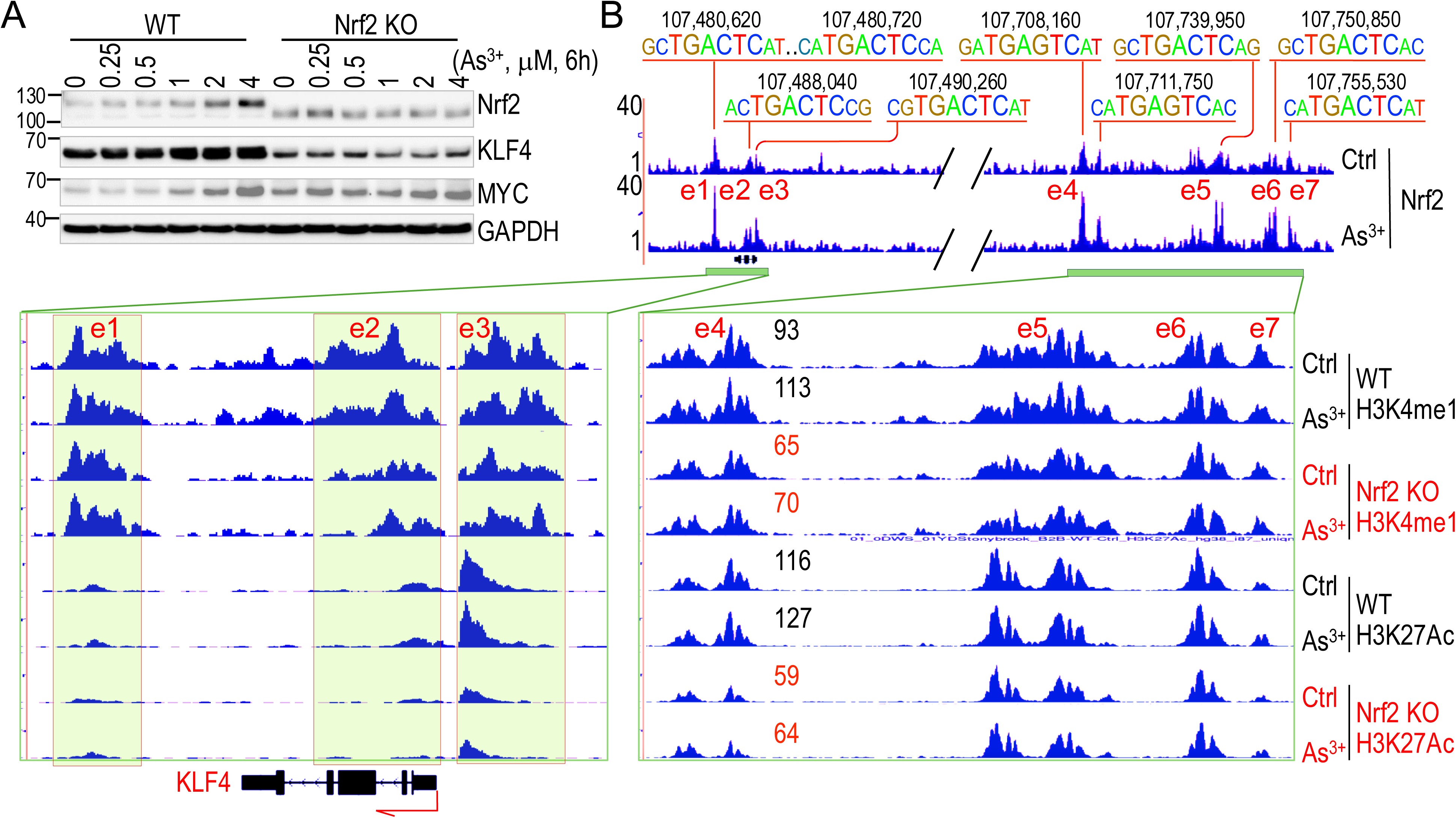
KLF4 is a transcriptional target of Nrf2. **A.** WT and Nrf2 KO cells treated with the indicated concentrations of As^3+^ for 6h followed by Western blotting analysis for the indicated proteins. **B.** As^3+^ enhanced Nrf2 binding to and enrichment of enhancer markers H3K4me1 and H3K27Ac in the KLF4 gene locus as determined by ChIP-seq. The ARE elements and the corresponding positions in the genome are shown at the top of the panel. Potential enhancers are designed as e1 to e7. Numbers in the bottom right panel show the relative degree of enrichment of the enhancer markers in WT and Nrf2 KO cells.

### Active transcription of KLF4 in the transformed WT cells

Similar to non-transformed cells, transformed WT cells displayed a dose-dependent induction of KLF4 and MYC proteins in response to As^3+^. However, in transformed Nrf2 KO cells, the induction of both KLF4 and MYC by As^3+^ was reduced (Fig. 3A). This differs from the non-transformed Nrf2 KO cells, which showed a reduction in KLF4 but not MYC (Fig. 2A). Compared to non-transformed WT cells, As^3+^-induced transformed WT cells showed a significantly elevated expression of genes associated with stem cells and CSCs, including TBX family members, KLF4, TCF4, SOX2, and several Wnt family genes (Fig. 3B). A transcription factor pathway analysis of 4008 protein coding genes, with an adjusted fold change of ≥ 3 when comparing transformed WT to non-transformed WT cells, identified the top transcription factors for these overexpressed genes as KLF11 and KLF4, followed by ZNF148, CACYBP, TFAP2A, and Nrf2 (Fig. 3C). KLF4 mRNA expression was significantly lower in transformed Nrf2 KO cells compared to transformed WT cells (Fig. 3D). It is well-established that the degree of H3K4me3 enrichment and peak breadth in ChIP-seq at the transcription start site (TSS) or promoter region correlates with the transcription status of genes. Genes with higher or broader H3K4me3 peaks are generally more actively transcribed [26–29]. In the Klf4 gene, two broad H3K4me3 peaks align with Nrf2 binding peaks: one at the upstream -510bp region of the TSS and another spanning from exon 1 to exon 3, peaking at the 770bp region (Fig. 3E). Transformed WT cells not only showed increased H3K4me3 enrichment but also had reduced levels of H3K9me3 and H3K27me3, two markers associated with transcriptional repression, at the Klf4 gene. Additionally, beyond the H3K4me3 peaks near the TSS, there are intergenic H3K4me3 peaks located 148kb upstream and 24kb downstream of the Klf4 gene. Both of these intergenic H3K4me3 peaks were enhanced by As^3+^ treatment in the transformed cells (Fig. 3E).

**Fig. 3.**
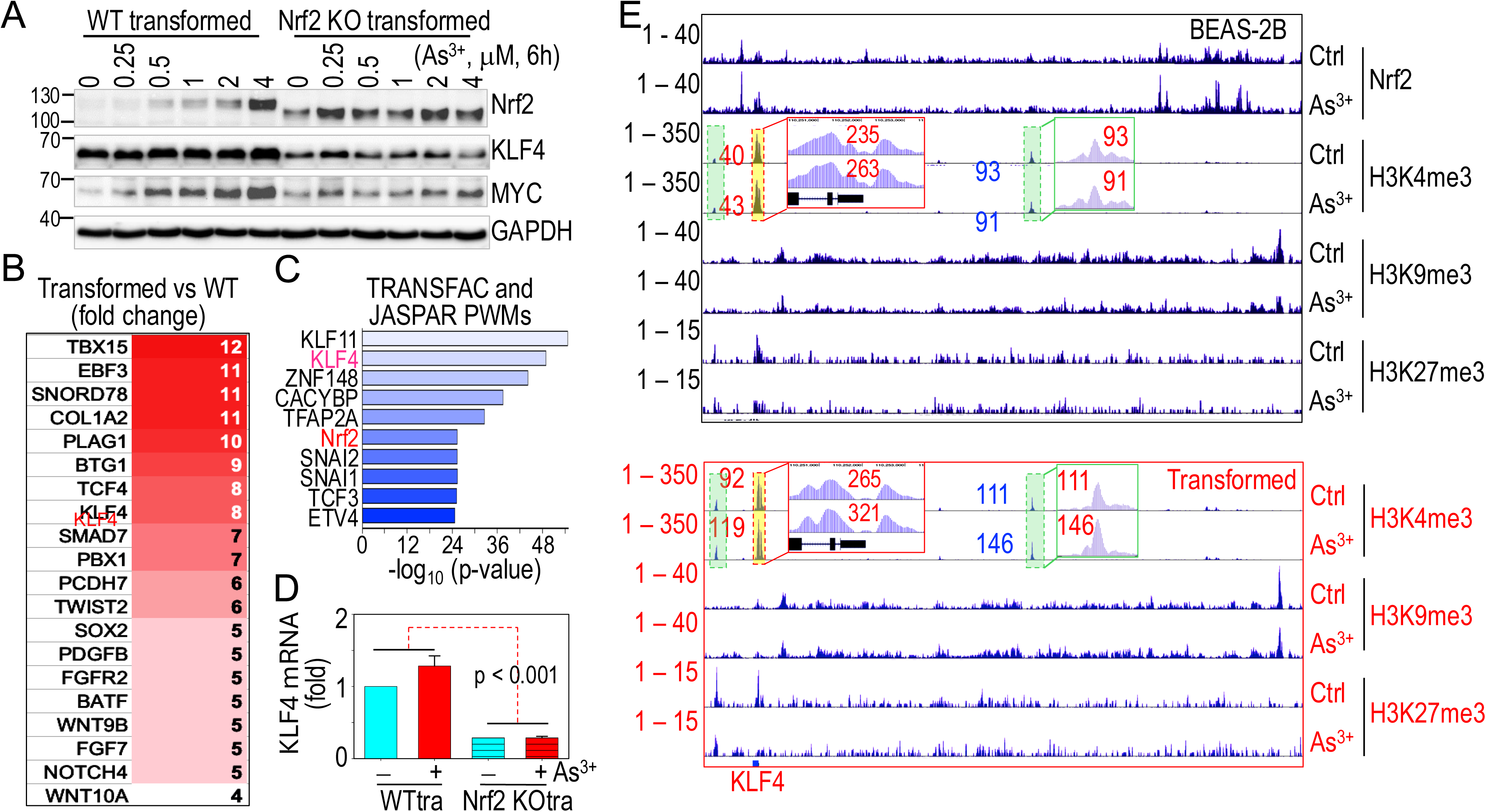
Elevated potential of KLF4 expression in transformed WT cells. **A.** Western blotting analysis for Nrf2 activation and expression of KLF4 and MYC in transformed WT and transformed Nrf2 KO cells treated with As^3+^. **B.** Expression levels of the genes associated with cancer stem-like cells (CSCs) in transformed WT cells relative to the non-transformed WT cells. **C.** Transcription factor pathway assay of the upregulated genes in transformed WT cells. **D.** Semi-quantitative RT-PCR showing KLF4 mRNA expression in control and As^3+^-treated transformed WT (WTtr) and transformed Nrf2 KO (Nrf2 KOtr) cells. **E.** ChIP-seq showing the histone methylation profiles of the indicated markers in Klf4 gene locus in non-transformed BEAS-2B cells (upper panel) and transformed WT cells (bottom panel). Spectrums of Nrf2 ChIP-seq in the control and As^3+^-treated non-transformed BEAS-2B cells were used at the top of the panel as position references of the indicated histone methylation markers. The H3K4me3 peaks at the TSS of KLF4 (highlighted in yellow) and up- and down-stream H3K4me3 peaks are highlighted in green. Depletion of Nrf2 using CRISPR-cas9 gene editing led to a reduction in both KLF4 expression and its chromatin binding, along with a decreased cell transformation induced by As^3+^

### Nrf2 participates in the enhancer remodeling

The abnormal activation of enhancers, particularly those linked to oncogenic and stemness genes, can drive self-sustaining transcriptional circuits in transformed or cancer cells. While overall levels of H3K4me1 and H3K27Ac did not differ significantly between WT and Nrf2 KO cells in ChIP-seq analysis, the loss of Nrf2 led to a significant reduction of these markers at the Klf4 gene (Fig. 2B). This suggests that Nrf2 is involved in the remodeling of Klf4 enhancers. Well-documented evidence shows that H3K4me1, catalyzed by KMT2C or KMT2D, marks the poised state of enhancers, and the recruitment of the acetyltransferase EP300 by H3K4me1 induces H3K27Ac to form fully active enhancers. To understand how Nrf2 knockout diminished H3K4me1 and H3K27Ac levels at the Klf4 locus, we first examined the impact of Nrf2 KO on the expression of methyltransferases responsible for H3K4me1, acetyltransferases for H3K27Ac, and genes associated with Compass complex, Cohesin complex, and BAF complex, which are involved in enhancer activities. As shown in Fig. 4A, KMT2D, a key methyltransferase for H3K4me1, was significantly decreased in Nrf2 KO cells. Additionally, acetyltransferases involved in H3K27Ac formation, including EP300, CREBBP, and ELP4, were slightly reduced following Nrf2 depletion. This finding was further supported by decreased levels of KMT2D and EP300 proteins in Nrf2 KO cells (Fig. 4B), indicating that Nrf2 is essential for the expression of these critical enhancer regulators. This observation aligns with the enrichment of Nrf2 and H3K4me3 at the promoter region of the KMT2D gene in As^3+^-treated and transformed cells. Notably, a conserved ARE site is present at the peak of Nrf2 enrichment (Fig. 4C), clearly indicating that KMT2D is a Nrf2-target gene involved in enhancer activity at the Klf4 gene locus and other gene loci.

**Fig. 4.**
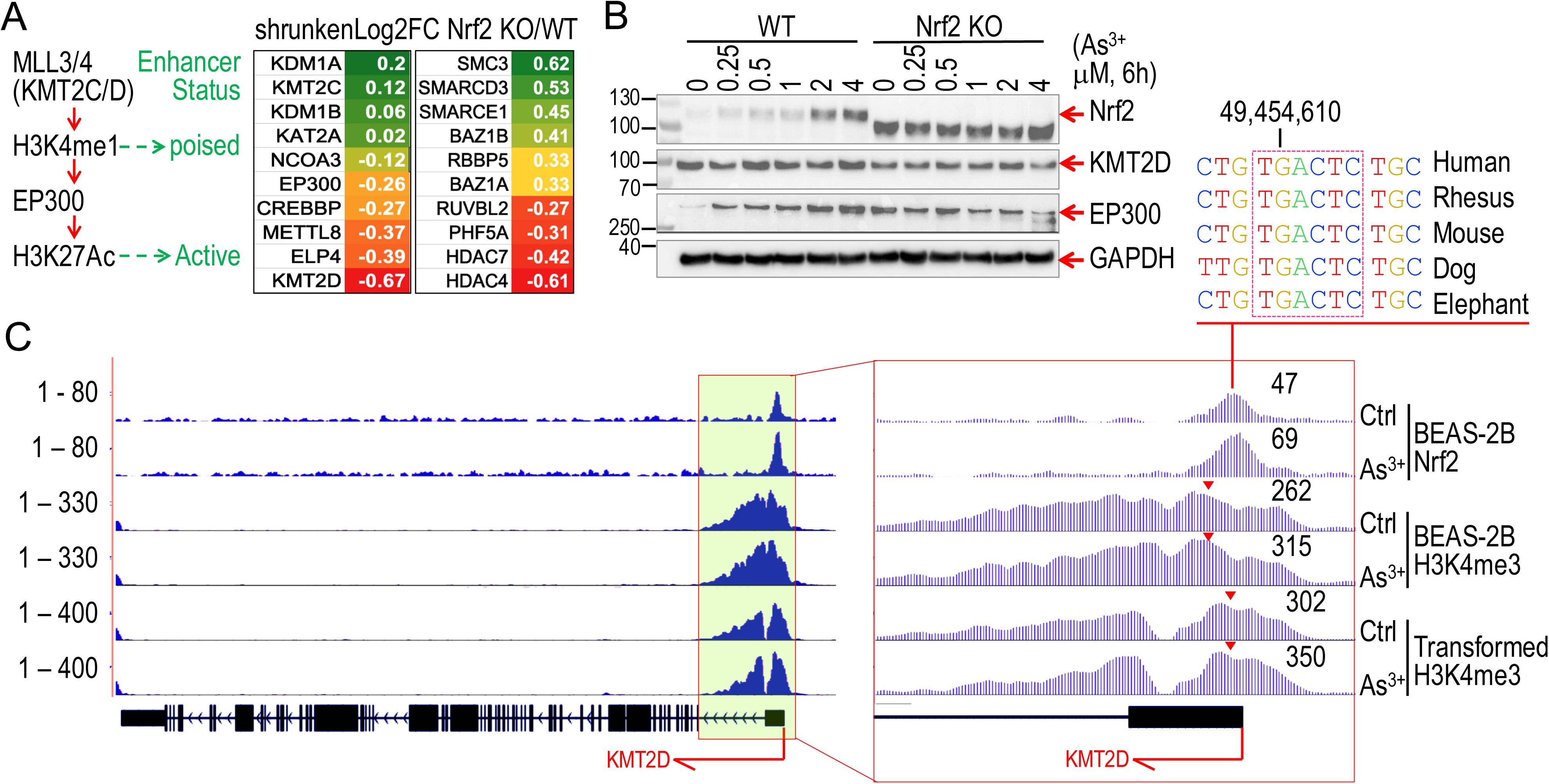
Nrf2 contributes to the establishment of enhancers. **A.** Diagram showing establishment of poised and active enhancers. Right panel indicates the expression levels of enhancer-related genes in Nrf2 KO cells relative to the WT cells. **B.** Knocking out Nrf2 reduces expression of KMT2D and EP300. **C.** ChIP-seq plot shows that As^3+^ enhances enrichment of Nrf2 and H3K4me3 at TSS of Kmt2d in the indicated cells. Conserved Nrf2 binding ARE element is indicated on the top of the panel.

### Ablation of Nrf2 diminishes the genome-wide binding of KLF4 in response to As^3+^

To further investigate the Nrf2 dependency of As^3+^-induced KLF4, we conducted global ChIP-seq analysis of KLF4 in both WT and Nrf2 KO cells. Consistent with the upregulation of KLF4 mRNA and protein, transient As^3+^ treatment led to a strong enrichment of KLF4 across the genome in WT cells, which was substantially inhibited in the Nrf2 KO cells (Fig. 5A). KLF4 binding peaks were predominantly located near TSS, with a moderate increase in KLF4 binding also observed upstream and downstream of genes (Fig. 5B). HOMER motif analysis revealed significant enrichment of six KLF family motifs, including KLF1, KLF3, KLF4, KLF5, KLF6, and EKLF, in all groups of cells. The top-three motifs identified among the KLF4 peaks in As^3+^-treated WT cells were KLF4, KLF1, and KLF6 (Fig. 5C). In contrast, SP2, KLF5, CG-repeat binding protein, AP1, KLF3, and KLF6 were overrepresented in control and As^3+^-treated Nrf2 KO cells. Under basal conditions, only a few minor KLF4 peaks were detectable across the chromosomes in WT cells, as illustrated by chromosome 3 (Fig. 5D). Multiple strong KLF4 clusters emerged in As^3+^-treated WT cells, whereas KLF4 binding peaks were nearly undetectable in the Nrf2 KO cells.

**Fig. 5.**
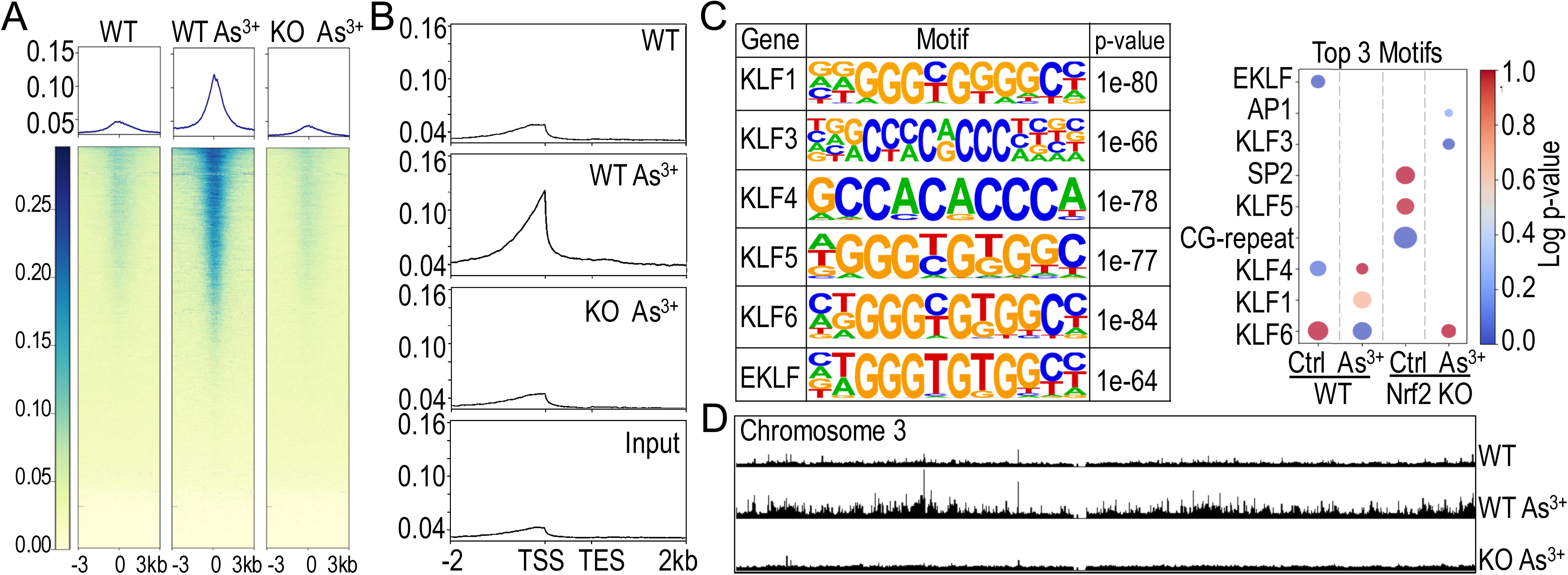
Knocking out Nrf2 blocks As^3+^-induced genome-wide KLF4 binding. **A.** ChIP-seq heatmaps of KLF4 binding in control and As^3+^-treated WT and Nrf2 KO cells. **B.** Enrichment distribution of KLF4 on gene loci. **C.** Top enriched KLF elements in the indicated cells with or without As^3+^ treatment. **D.** Screenshot of KLF4 binding as determined by ChIP-seq on the entire chromosome 3 of the indicated cells.

### Unique characteristics of KLF4-regulated genes

KLF4 is best known for its role in maintaining self-renewal in stem cells and iPSCs. However, its transcriptional targets in differentiated cells remain unclear or controversial. In WT cells, KLF4 ChIP-seq data revealed that As^3+^-treatment increased KLF4 enrichment on several thousands of genes, a process that can be reversed by Nrf2 deficiency (Fig. 6A). Analysis of genes with enhanced KLF4 enrichment in response to As^3+^ revealed a preference for KLF4 binding to genes with longer 3’-UTR and lower GC content in their sequences (Fig. 6B, top panels). Gene type distribution profiling showed that KLF4 primarily binds to protein-coding genes, with a reduced affinity for long non-coding RNAs, pseudogenes, and other non-coding RNAs (Fig. 6B, bottom panel). Although only about 30% of KLF4-enriched genes showed increased expression in As^3+^-induced transformed cells, 16 out of the top 17 KLF4-enriched genes, with the exception of HMGB1, were overexpressed in these transformed cells (Fig. 6C). Pathway analysis of As^3+^-induced KLF4-enriched genes indicated an overrepresentation of transcription factors such as STAT3, OVOL1, GATA2, JUN, SOX2, and Nrf2, as well as functional pathways related to gene expression, cell migration, and EMT (Fig. 6D). Interestingly, several Nrf2-regulated genes, including Nrf2 itself, MYC, CD44, KMT2D, and OSER1, exhibited As^3+^-induced KLF4 binding (Fig. 6E). Additionally, As^3+^ triggered significant KLF4 enrichment on WNT10A and specific immune checkpoint genes such as NECTIN2 (CD112), PDCD1 (PD-1), and CD274 (PD-L1), whose expression is Nrf2 dependent, despite the absence of detectable Nrf2 binding to these genes in ChIP-seq analysis (Fig. 6E and data not shown).

**Fig. 6.**
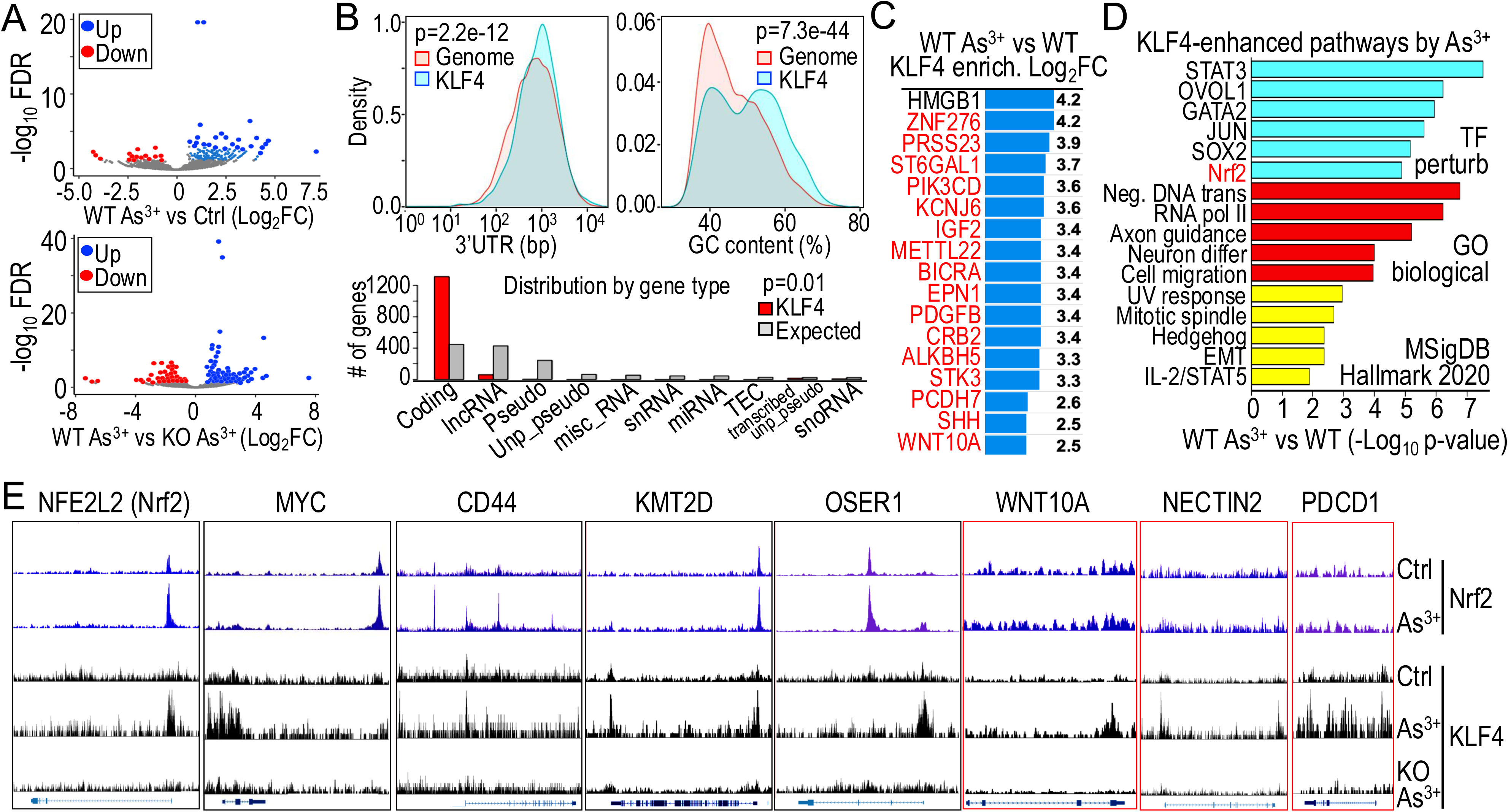
Identifying KLF4 target genes through ChIP-seq. **A.** Volcano plots of genes regulated by KLF4 As^3+^-treated cells vs control cells (up panel), and As^3+^-treated WT cells vs As^3+^-treated Nrf2 KO cells. **B.** Characteristics of the KLF4-regulated genes in response to As^3+^. **C.** Top KLF4-enriched genes induced by As^3+^. Genes marked in red are upregulated in As^3+^-induced transformed cells. **D.** Pathway analysis of KLF4-regulated genes in response to As^3+^. **E.** KLF4 binds to Nrf2-target genes with or without Nrf2-binding peaks in Nrf2 ChIP-seq.

### Concerted regulation of Nrf2 and KLF4 on enhancers

At the chromosome level, we observed a tendence for co-localization of As^3+^-enhanced Nrf2 and KLF4 with elevated levels of H3K4me1 and H3K27Ac (Fig. 7A). This pattern is also evident at the individual gene level, as exampled by EGFR. The As^3+^-induced Nrf2 peak overlaps with the KLF4 peak within the downstream of intron 1, where high levels of enhancer markers H3K4me1 and H3K27Ac are present. Notably, enhancer marker levels, especially H3K27Ac, were lower when either Nrf2 (highlighted in yellow) or KLF4 (highlighted in bright pink) was enhanced individually (Fig. 7A, right). Further support for this observation comes from the correlation between Nrf2, KLF4, and enhancer markers on the KLF4 gene itself. In addition to increased Nrf2 binding in downstream, immediately upstream, and within the gene body, As^3+^ treatment also elevated KLF4 binding at the immediate upstream region and the gene body, spanning exon 1 to exon 3, where more than 15 KLF4-binding elements, some of which are clustered together, were identified (Fig. 7B). The density of KLF4 elements is proportionally correlates with the height of KLF4 peaks in ChIP-seq, with the highest KLF4 clusters located adjacent to a strong Nrf2 peak containing a conserved ARE element, and roughly aligned with the highest enhancer markers, particularly H3K27Ac. Interestingly, no detectable KLF4 binding peaks and only a marginal H3K27Ac peak were observed in the downstream of the KLF4 gene, despite the presence of a strong As^3+^-induced Nrf2 peak. These findings clearly indicate that following Nrf2 activation, KLF4 is highly capable of self-promoting its own gene expression, and the co-localization or close proximity of Nrf2 and KLF4 facilitates the establishment of active enhancers.

**Fig. 7.**
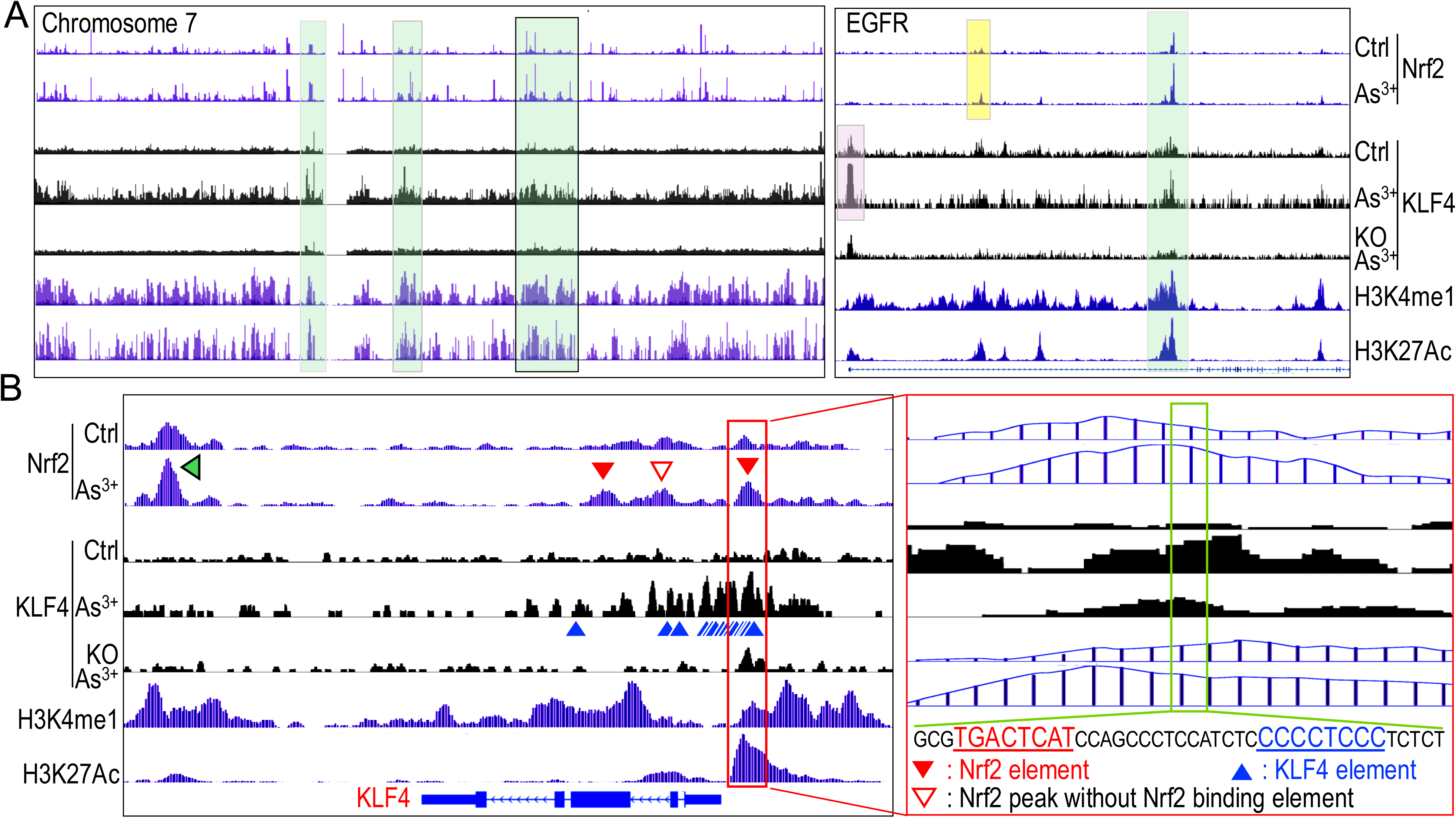
Co-occupancy of Nrf2 and KLF4 facilitates enhancer institution. **A.** Representative distribution and co-localization of Nrf2, KLF4, H3K4me1, and H3K27Ac in chromosome 7 (left) and on the gene body of Egfr (right). Regions highlighted with green boxes show co-occupancy of Nrf2 and KLF4 with the active enhancer markers H3K4me1 and H3K27Ac. **B.** Self-amplification of KLF4 through enhanced KLF4 binding to the gene body and upstream of the Klf4 gene. Red box shows co-occupancy of Nrf2 and KLF4 with the active enhancer. Blue triangles indicate KLF4 binding elements; red filled triangles depict Nrf2 peaks contain ARE element; red unfilled triangle indicates Nrf2 peak without typical ARE element; green triangle points to the strongest Nrf2 enrichment peak located in a region without KLF4 binding peaks nor active enhancer marker H3K27Ac in the proximal downstream of Klf4 gene.

## Discussion

We present previously undocumented evidence demonstrating the role of Nrf2 in promoting the expression of the stemness factor KLF4 through the coordinated regulation of both enhancers and KLF4 itself, further reinforcing the pro-carcinogenic nature of Nrf2 in cancer cells and cancer stem-like cells (CSCs). The use of CRISPR-cas9 gene editing to deplete Nrf2 resulted in a decrease in KLF4 expression and its chromatin binding, as well as a reduction in cell transformation induced by As^3+^. ChIP-seq analysis revealed multiple Nrf2 enrichment peaks, particularly under As^3+^-induced stress conditions, at the KLF4 gene locus. Most of these Nrf2 peaks contain conserved ARE elements, confirming the authenticity of Nrf2 regulation on KLF4. Beyond direct transcriptional regulation, Nrf2 may also promote KLF4 expression by facilitating the establishment of enhancers at the Klf4 locus and other loci, as evidenced by the fact that expression of KMT2D, a methyltransferase for H3K4me1, is also partially Nrf2 dependent. Intriguingly, Nrf2 collaborates with its target gene KLF4 in the formation of active enhancers. Furthermore, KLF4 contributes to its own expression by binding to multiple KLF4 elements within the gene body and upstream of the KLF4 gene.

As one of the original four Yamanaka factors identified for reprogramming iPSCs, KLF4 has been extensively studied in the context of self-renewal and lineage development in pluripotent stem cells and progenitor cells. However, its role as an oncogenic factor or tumor suppressor during carcinogenesis and tumor progression remains unclear. One of the earliest findings supporting KLF4’s oncogenic role is the attenuating effect of Klf4 ablation on mutant Kras^G12D^-induced pancreatic intraepithelial neoplasia in a mouse model of pancreatic cancer [30]. It is believed that stress signal-induced KLF4 is responsible for driving the acinar-to-ductal transition, which is crucial for the formation of acinar-to-ductal metaplasia (ADM) and pancreatic intraepithelial neoplasia (PanIN), two common pre-malignant lesions of pancreatic cancer. KLF4’s oncogenic role is further demonstrated in the metastatic migration and invasion of breast cancer cells where KLF4 is overexpressed [31]. Similarly, KLF4 has been shown to be critical for metastasis and stemness of the osteosarcoma cells in response to chemotherapy [32]. In human lung cancer, KLF4 expression varies depending on the cancer type and stage. Increased KLF4 expression is observed in small-cell lung cancer but not non-small-cell lung cancer, with higher rate of expression noted in stages II, III and IV [33]. Intriguingly, some studies have also demonstrated a tumor suppressor-like role for KLF4 in experimental cancer models and human cancers. Overexpressing KLF4 in murine prostate stem cells impedes malignant transformation, whereas KLF4 loss induces the molecular features characteristic of aggressive tumors [34]. These paradoxical roles of KLF4 in cancer may depend on the specific cellular context and the relative abundance of KLF4 and other oncogenic or tumor suppressive proteins. For example, KLF4 may act as a tumor suppressor in normal cells by inducing the expression of the tumor suppressor p21CIP1 (CDKN1A), but as an oncogene in transformed or cancer cells by antagonizing p53 function [35]. The complexity of KLF4’s role in cancer is further underscored by the functional and structural similarities within the KLF protein family, which consists of 18 members and can be induced by a wide range of stress signals [36, 37].

We and others have previously reported that Nrf2 acts as a transcriptional regulator for KLF4 in human cells [10, 38]. However, the multiple Nrf2 binding sites, or ARE elements, within the Klf4 gene locus that showed strong Nrf2 enrichment in response to As^3+^ have not been reported before. Particularly intriguing is the presence of Nrf2-binding clusters overlapping with the enhancer markers H3K4me1 and H3K27Ac located 218 kb upstream of the Klf4 gene. In HeLa cells, this upstream region appears to be crucial for forming a topologically associating domain (TAD) between the upstream distal enhancer e7 (Fig. 2) and the Klf4 promoter, or downstream proximal enhancer e1 and distal enhancer e7, facilitating Klf4 transcription (data not shown). Nrf2 knockout led to reduced KLF4 expression, along with a concomitant decrease in enhancer markers overlapping with Nrf2 clusters, suggesting that, in addition to direct transcription regulation, Nrf2 can promote KLF4 expression through enhancer activation and TAD formation. Indeed, our data indicate that Nrf2 is a key regulator of KMT2D, a methyltransferase responsible for establishing enhancers via H3K4me1.

Despite decades of extensive research, the role of Nrf2 in regulating enhancers or super-enhancers remains largely unexplored. Early studies demonstrated that the Neh4 and Neh5 domains of Nrf2 physically interact with CREBBP (CBP/p300) [39, 40], an acetyltransferase that catalyzes the formation of H3K27Ac to activate enhancers [41]. This suggests that Nrf2 binding to ARE elements could recruit CREBBP, facilitating enhancer activation. In contrast, KLF4’s association with enhancers is well-established, particularly in the reprogramming of iPSCs. Emerging evidence indicates that KLF4 can orchestrate the topological reorganization of enhancer-based TADs, or genome-wide enhancer rewiring [42], in addition to its ability to recruit and physically interact with CREBBP [43]. However, in differentiated or cancer cells, KLF4 alone may be insufficient for CREBBP recruitment, enhancer activation, or chromatin architectural reorganization, suggesting that additional architectural factors or co-regulators are necessary. In this context, Nrf2 is a promising candidate to cooperate with KLF4 in fully establishing and activating enhancer hubs and their associated topological assemblies. The finding in this report that co-occupied regions of Nrf2 and KLF4 exhibit features of active enhancers provides strong support for this hypothesis.

In summary, we present evidence that Nrf2 not only directly promotes KLF4 expression but also supports KLF4’s chromatin binding and architectural function in enhancer activation and TAD reorganization. These findings provide a mechanistically explanation for why gain-of-function mutations in Nrf2 are frequently linked to aggressive tumors, drug resistance, recurrence, and metastasis. As such, targeting Nrf2 represents a promising strategy for cancer therapy, potentially limiting the generation and expansion of CSCs responsible for metastasis and tumor recurrence. Our understanding of the roles Nrf2 and KLF4 play in enhancer function and chromatin architecture is still in its early stages. Further investigation into their interactions and oncogenic roles is urgently needed.

